# The Effects of Physical and Mental Fatigue on Time Perception

**DOI:** 10.1101/2023.12.05.570265

**Authors:** Reza Goudini, Ali Zahiri, Shahab Alizaheh, Benjamin Drury, Saman Hadjizadeh Anvar, Abdolhamid Daneshjoo, David G Behm

## Abstract

The subjective perception of time holds a foundational significance within the realm of human psychology and our conceptualizations of reality and how we elucidate the chronological progression of events within our lives. While there have been some studies examining the effects of exercise on time perception during the exercise period, there are no studies investigating the effects of fatiguing exercise on time perception after the exercise intervention. This study aimed to investigate the effects of physical and mental fatigue on time estimates over 30-seconds (5-, 10-, 20-, and 30-seconds) immediately after exercise and 6-minutes after the post-test. Seventeen healthy and recreationally active volunteers (14 males, 3 females) were subjected to three conditions: physical fatigue, mental fatigue, and control. All participants completed a familiarization and three 30-minute experimental conditions (control, physical fatigue (cycling at 65% peak power output), and mental fatigue (Stroop task for 1100 trials) on separate days. Heart rate and body temperature were recorded at the pre-test, start, 5-, 10-, 20-, 30-seconds of the interventions, post-test, and 6-min follow-up. Rating of perceived exertion (RPE) was recorded four times during the intervention. Time perception was measured prospectively (at 5-, 10-, 20-, and 30-seconds) at the pre-test, post-test, and 6-minute follow-up. Physical fatigue significantly (p=0.001) underestimated time compared to mental fatigue and control conditions at the post-test and follow-up, with no significant differences between mental fatigue and control conditions. Heart rate, body temperature, and RPE were significantly (all p=0.001) higher with physical fatigue compared to mental fatigue and control conditions during the intervention and at the post-test. This study demonstrated that cycling-induced fatigue led to time underestimation compared to mental fatigue and control conditions. It is crucial to consider that physical fatigue has the potential to lengthen an individual’s perception of time estimates in sports or work environments.

## Introduction

People have been fascinated by time for centuries; however, philosophers and scientists from ancient to modern times have yet to fully agree on its definition and qualities (Bunnag, 2019). The concept of time is one of the experiences that are essential for how we experience the world (Wittmann, 2009). Our behavioral and cognitive systems depend heavily on duration perception, which allows us to interact with the outside world (Jia and Feng, 2020). An accurate perception of time is an indispensable part of many time-constrained sports (i.e., North American football, basketball, figure skating, and others) and work environments (Behm and Carter, 2020). It is well known that our subjective perception of time can be manipulated and distorted under certain circumstances (Eagleman, 2005); however, little is known about how physical and mental fatigue affects how people perceive time.

There are two prominent theories pertaining to time perception: the Pacemaker Accumulator Model (PAM), alternatively referred to as the Scalar Expectancy Theory (SET) (Gibbon, Church and Meck, 1984), and the Striatal Beat Frequency Model (SB-FM) (Meck, 1983; Meck and Church, 1983; Grondin, 2010). Both theoretical frameworks elucidate that time perception is significantly impacted by arousal (Grondin, 2010; Allman and Meck, 2012). The scalar expectancy theory uses a clock, memory, and decision stages to divide the temporal processing system. The SB-FM not only discusses the timing behaviors, but also identifies which neural regions of the brain are involved (Merchant and Meck, 2013). The model suggests that time estimation is determined by dopamine and glutamate activity near the substantia nigra and ventral tegmental area. The timing process starts with striatal spiny neurons that monitor activation patterns in the cortex’s oscillatory neurons, which are controlled by glutamate action (Meck, 2005). The oscillating neurons synchronize when an interval starts, and the spiny neurons are reset by phasic dopaminergic input. A dopamine pulse is released when the target duration is attained, strengthening the synapses that are active in the striatum (Meck, 2005). Time is perceived in the mind according to the oscillatory activity rate. Once the same signal duration has been timed once more, neostriatal GABAergic spiny neurons compare the current activation pattern to the stored pattern to determine when the duration has been reached; when they match, spiny neurons fire to indicate that the period has passed (Matell and Meck, 2004; Meck, 2005; Merchant et al., 2013).

Arousal associated with physical and mental activity can increase heart rate, muscle activation (e.g., motor unit recruitment and firing frequency), thermoregulation, and other physiological or external signals, that have the potential to alter time perception (Graham et al., 2023). The increased activity gives rise to additional events within the temporal processing system, leading to an accelerated perception of time in response to higher-intensity contractions. Given the cerebellum’s involvement in both movement and temporal processing (Ivry and Diener, 1988), exercise-induced arousal may exert more influence on time perception compared to other forms of arousal. The heightened demands in terms of frequency of events during sensory afferent processing may also play a role in impacting time perception (Graham et al., 2023).

As a psychobiological condition, mental fatigue (e.g., difficulty in maintaining focus, attention, cortical excitability) results from extended periods of demanding cognitive activity (Job and Dalziel, 2000). It has been shown that mental fatigue increases prefrontal cortex activation, cerebral perfusion, and somatomotor activation. This increase in blood flow to the prefrontal cortex may result from additional neural activation required to produce efferent motor commands, which may be one of the reasons why central fatigue can have an impact on physical performance (Mehta and Parasuraman, 2014; Nobrega et al., 2014). Previous studies have shown that the way in which central fatigue manifests depends on the specific task, with continuous low-moderate intensity exercise frequently leading to greater central fatigue (Place and Westerblad, 2009; Kennedy et al., 2013; Krüger et al., 2019; Iannetta et al., 2022). After finishing a mentally exhausting task, an approach is to look at behavioral performance deficiencies on a subsequent task (Helm, 2021). The best explanation for behavioral changes during such tasks is a reduction in top-down processing, which results in an inability to focus and meet task demands.

Exercising for a long time in a hot environment impairs physical and mental performance (González-Alonso et al., 1999). Timing behaviour is sensitive to changes in body temperature; hence it has been suggested that there is a temperature-sensitive time mechanism (Tamm et al., 2015). Some studies claim that a rise in temperature causes time to hasten, but others contend that this effect only happens once a certain threshold of perceived fatigue has been reached (Tamm et al., 2014, 2015). Exercise encompasses a wide range of forms, with differences in intensity, duration, and movement types. However, time perception research often overlooks the significance of contraction types, such as dynamic contractions. Additionally, the impact of varying fatigue protocol on time perception remains relatively unexplored, representing an understudied factor that could potentially affect time perception positively or negatively.

Since no studies have compared physical and mental fatigue on time perception, the objective of this study was to compare the effects of physical (exhaustive cycling exercise protocol) and mental fatigue (Stroop task test) on the time perception. It was hypothesized that physical and mental fatigue protocol will lead to an underestimation of the time.

## Methods

### Participants

An “a priori” statistical power analysis (software package, G * Power 3.1.9.7) was conducted based on the time perception of related studies (Tonelli, Lunghi and Gori, 2022) to achieve an alpha of 0.05, an effect size of 0.4, and a statistical power of 0.8 using the F-test family. The analysis indicated that between 12-14 participants per condition was sufficient to achieve adequate statistical power. Seventeen (17) healthy and recreationally active participants took part voluntarily in this study. Exclusion criteria included: participants who have neurological conditions, knee injuries, presence of medical issues that prevent high-intensity exercise, or injuries to the quadriceps muscles that could affect pedaling. Inclusion criteria included that participants need to be healthy, and recreationally active. The recruitment period for participants began January 16, 2023 and was completed by September 8, 2023.

**Table 1:** Participant anthropometrics.

| Participants | Age (years) | Mass (kg) | Height (cm) |
| --- | --- | --- | --- |
| Male (n=14) | 28.57 ± 4.92 | 80.46 ± 11.22 | 175.71 ± 2.65 |
| Female (n=3) | 24 ± 2.64 | 69.06 ± 10.96 | 157.33 ± 2.30 |

Prior to their lab visit, participants were given instructions to avoid intense activity (24 hours prior to participating) and to stop drinking alcohol, smoking, and using caffeine (12 hours). Each participant completed the physical activity readiness questionnaire plus (PAR-Q+ 2020), read and signed the informed consent form prior to testing and after a brief explanation of the study and the experiment’s procedures. During their first visit to the lab, every participant became familiar with all psychological measurements. The Institutional Health Research Ethics Board (ICEHR #20231533-HK) gave its approval for this study, which was carried out in accordance with the most recent version of the Helsinki Declaration.

### Experimental design

The effects of 30 minutes of physical and mental fatigue on time perception were investigated using a randomized crossover study design. The participants became familiar with a basic orientation to the testing procedures and equipment during the initial familiarization session and, they performed an incremental cycling test using Velotron ergometer (Velotron RacerMate, Seattle, USA) to determine their peak power output (PPO). The participants then came to the lab for three distinct testing sessions, physical fatigue, mental fatigue, or control. Each session was randomized and separated by at least 48 hours (Figure 1).

**Figure 1:**
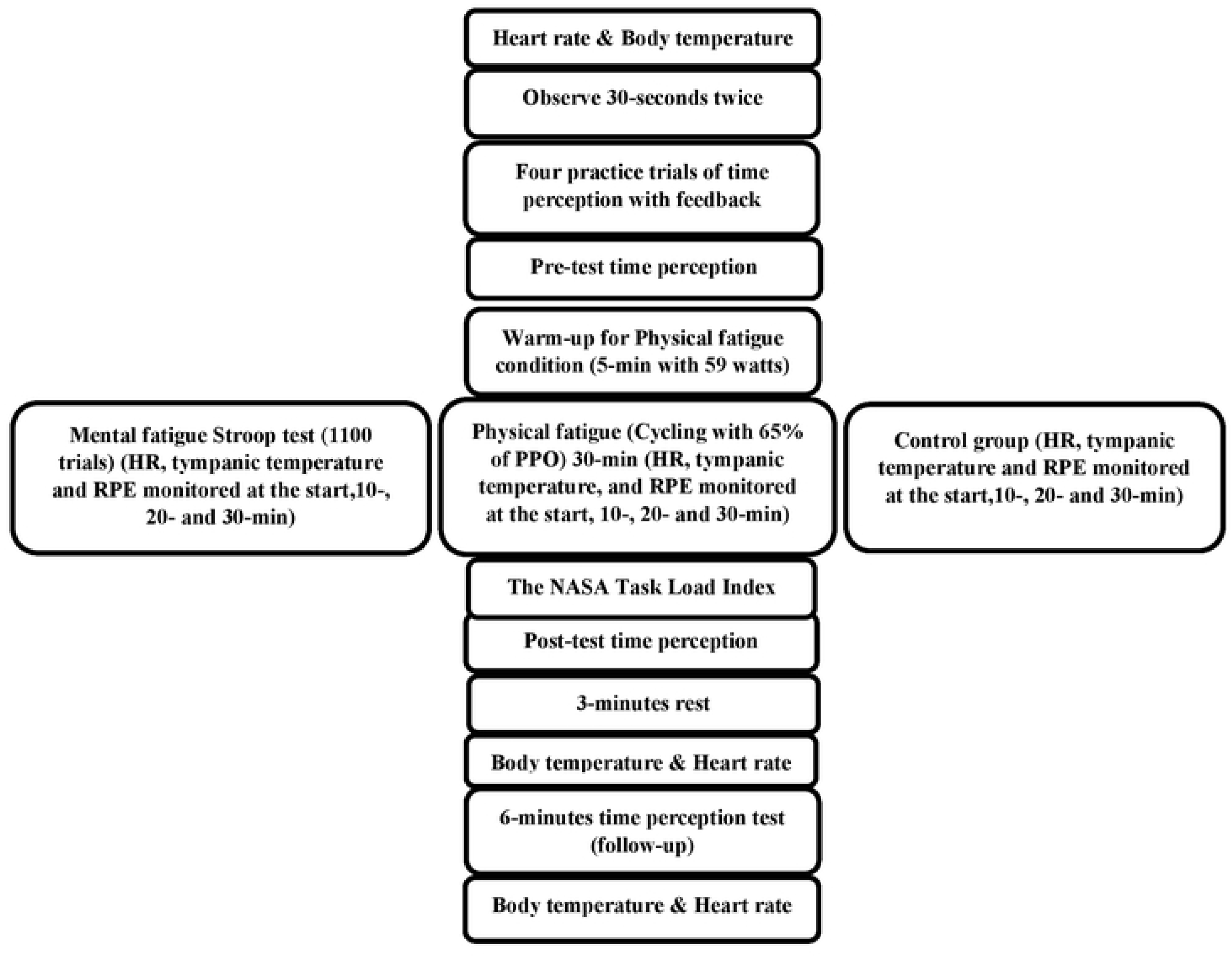
Experimental Design: PPO: peak power output, RPE: rating of perceived exertion

**Figure 2.**
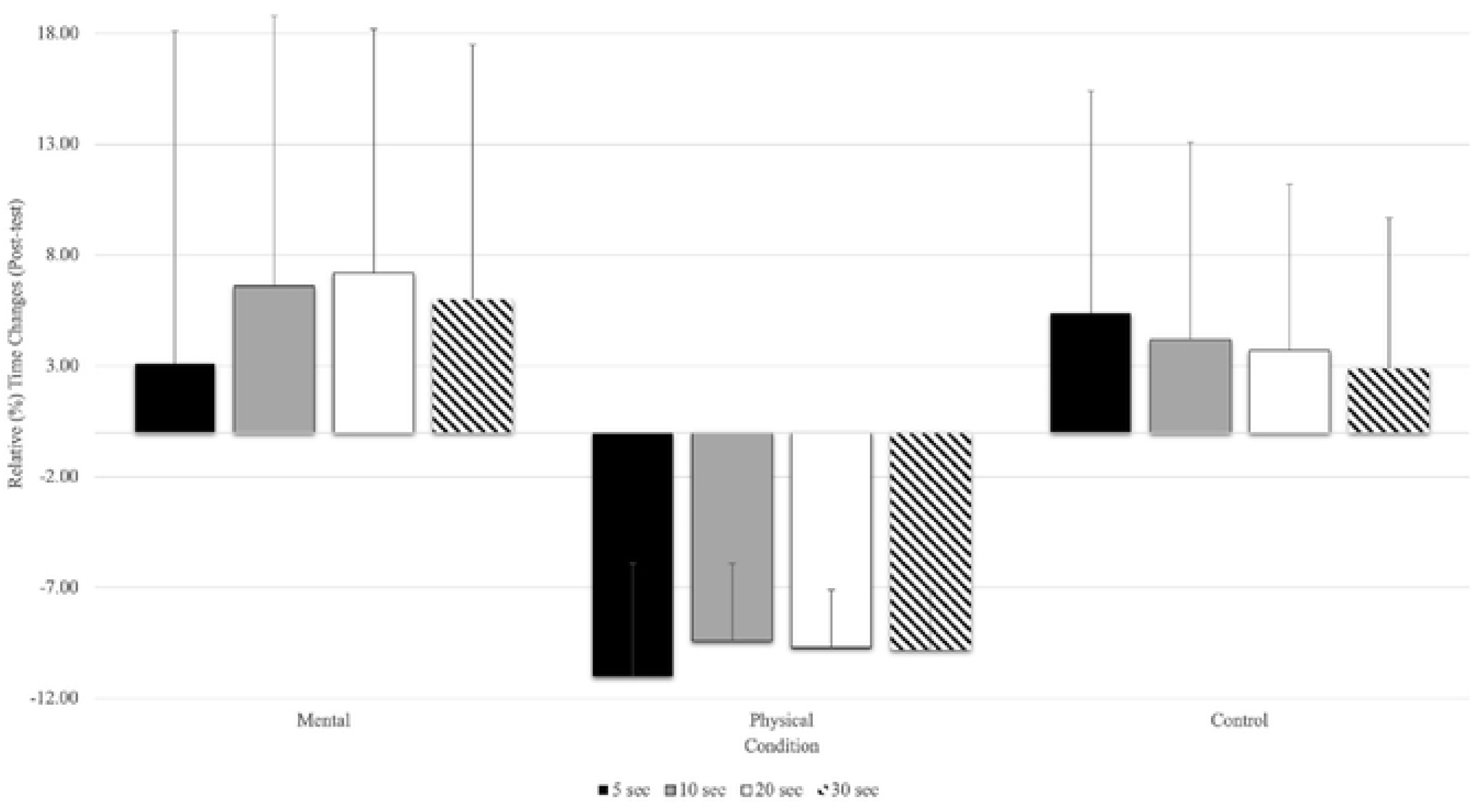
Relative (%) time change for the time estimates at the post-test (mean % ± SD). Only the physical fatigue condition significantly (asterisk represents p<0.01) underestimated time.

**Figure 3.**
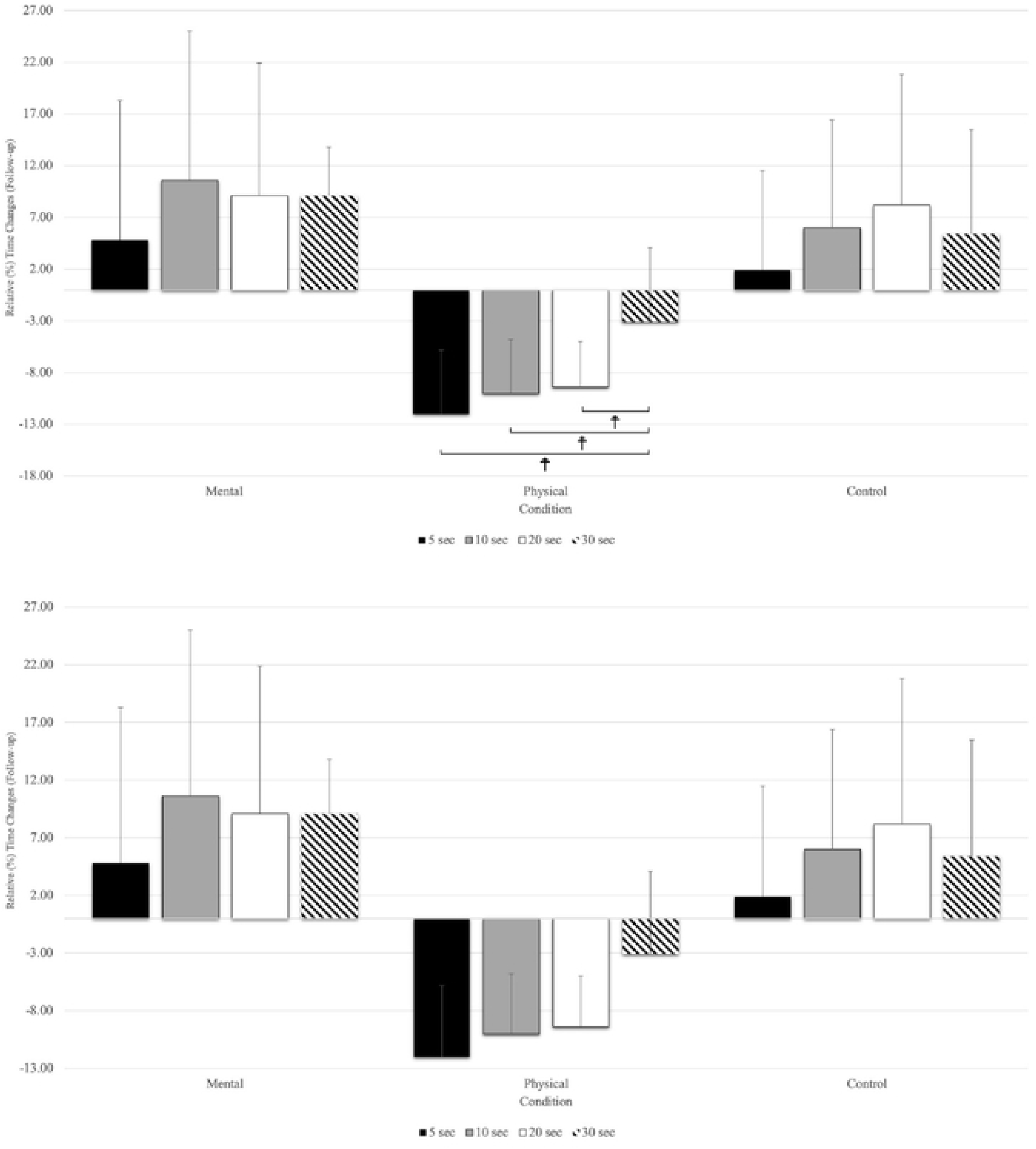
Relative (%) time change for the time estimates at the follow-up (mean % ± SD). Only the physical fatigue condition significantly (main effect for condition: p<0.001) underestimated the time. ☨ indicates p < 0.001 compared to the other time estimates.

### Measures

Prior to the intervention, participants watched a digital clock count to 30-seconds, twice, followed by four trials of time estimate practice of 30-seconds duration (estimate 5-, 10-, 20- and 30-seconds) with feedback. Then, for the pre-test data collection participants sat in a chair to estimate the time intervals of 5-, 10-, 20-, and 30-seconds, six times without feedback. We chose to execute the intervals six times as individuals can ingrain this into memory (as stated by the Scalar expectancy theory). Thirty seconds was chosen as this approximate time restriction is common in a number of sports including basketball, tennis, North American football, and others. This procedure has also been used successfully in prior experiments conducted in this lab with intraclass correlation coefficients (ICC) of 0.75-0.85 (Gardner et al., 2023; Graham et al., 2023). In the present study, a high degree of reliability was found between time perception measurements, with an ICC of 0.802 with a 95% confidence interval from .628 to .916 (F^(16,176)^ = 5.058, p<0.001). With the six 30-second time estimate attempts for each testing time, the mean scores were analyzed, for the pre-test, immediately post-test, and 6-minute follow up. Since six 30-second time estimates equals 3-minutes, it was decided to be consistent and permit a 3-minute recovery before the next 3-minute testing period (6 minutes in total). To estimate time, a hand dynamometer (custom built design) connected to a BioPac AcqKnowledge data acquisition system (Massachusetts, USA) was used.

Heart rate was monitored (T31, Polar, Kempele, Finland), tympanic temperature (IRT6520CA ThermoScan, Braun, Germany) and rating of perceived exertion (RPE) (Borg, 1998) were recorded in the pre-test, during the experimental protocols (start, 10-, 20-, and 30-minutes), and at the post-test. The heart rate monitor was fixed using an elastic belt secured around the participant’s sternum. Tympanic temperature was acquired with a thermometer’s probe, fitted with a disposable plastic covering, which was gently inserted into the right ear canal.

#### Rate of Perceived Exertion (RPE)

The RPE Borg Scale (Borg, 1998) was used as a tool for assessing the intensity of participants’ activity during the intervention, utilizing a graduated scale ranging from 6 to 20. Throughout the physical, mental fatigue, and control conditions, participants were prompted to provide their RPE ratings. The main aim of using the RPE Borg Scale was to gain valuable insights into whether participants were engaging in the prescribed activity at the desired intensity (Graham et al., 2023).

### Protocol

Prior to the pre-test, during the intervention (start, 10-, 20-, and 30-minutes), after the time perception test, and after the 6-minutes time perception test (follow-up) participants’ body temperatures and heart rates were recorded. The rating of perceived exertion was also recorded at the start, 10-, 20-, and 30-minutes during the intervention for the three conditions.

#### Maximal incremental cycling test protocol

The maximum cycling exercise protocol was used to determine the maximum wattage (W_max_) for the incremental test on a cycle ergometer (Velotron RacerMate, Seattle, USA). Each participant’s ideal seat height on the cycle ergometer was determined, recorded, and used for the following sessions. Participants warmed up with 59 watts with the RPM of 70 for 5-minutes and then participants began cycling at 80 watts for 3-minutes with RPM of 70, then raised their resistance by 40 watts every 3-minutes until they reached exhaustion (a cadence of less than 60 RPM for more than 5-seconds despite intense verbal encouragement). The researcher verbally encouraged participants during the test to perform a true all-out effort. The W_max_ (i.e., peak power output (PPO)) was calculated with the formula: W_max_= W_out_ + (t/180) × 40 [W_out_: workload of the last completed stage; t: time (seconds) in the final stage] (Barzegarpoor et al., 2020).

### Exhaustive cycling exercise protocol

Following the orientation practice time estimate sessions (two observations of a clock showing 30-seconds followed by four estimates of 30-seconds with feedback) and pre-tests, participants warmed up on the cycle ergometer for 5-minutes (with 59 watts) and then cycled at 65% PPO for 30-minutes. On the cycle ergometer, participants’ positions were adjusted to replicate their maximum cycling exercise. After the exhaustive cycling exercise protocol, participants filled out the NASA Task Load Index and then they were tested with six-time estimate trials immediately as well as six minutes after the immediate post-test time estimates.

### Stroop Task

The Stroop Colour-word test, a widely used neuropsychological test, measures a subject’s capacity to suppress cognitive interference, which happens when the processing of one stimulus attribute interferes with the concurrent processing of another (Stroop, 1935). It has been demonstrated that the Stroop task, which demands prolonged attention and response inhibition, induces a state of mental fatigue (Smith et al., 2016). The participant was asked to identify the colour of the word without regard for its actual meaning. Fifty percent (50%) of the trials were congruent (matched word and color), whereas 50% were incongruent, according to a pseudo-random sequence that was used to govern the trials (with all incongruent word-color combinations). The participants were then instructed to push the key on the keyboard that matches the color of the text that is displayed on the screen. The computer screen was 33 cm, and all participants used this laptop to observe 30-second time estimate and Stroop tasks throughout the study. For 1000 ms, each word appeared on the screen in font size 34, and then the screen remained blank before the next word appeared (Barzegarpoor et al., 2020). In this investigation, we conducted a total of 1100 trials to induce mental fatigue, requiring an approximate duration of 30 minutes for its completion. Slimani et al (2018), in their study showed that performing the Stroop task for 30 min successfully induced mental fatigue. For online data collection (Stroop task test), the software PsyToolkit was used (Stoet, 2010, 2017).

### NASA-TLX

The NASA-TLX was implemented for all three conditions. It is a tool for measuring mental workload that aims to record workers’ subjective perceptions of complex socio-technical systems that involve humans and machines. Due to its multidimensional nature and ease of administration, the NASA-TLX is perhaps the most commonly used mental workload scale (Colligan et al., 2015). NASA-TLX has six subscales that measure mental demand, physical demand, temporal demand, performance, effort, and level of frustration (Hart and Staveland, 1988). Following the implementation of interventions in each experimental condition, participants manually completed the NASA-TLX questionnaire. Participants were required to rate each item using a scale consisting of 20 equidistant intervals delineated by bipolar descriptors (e.g., high/low). Subsequently, the computed score was scaled by a factor of 5, yielding a resultant score ranging from 0 to 100 for each of the subscales (Barzegarpoor et al., 2020).

### Control Condition

The control condition executed the six trials (pre-test), watched a documentary film “When We Left Earth: The NASA Missions – Episode 6: A Home in Space” (Discovery Channel, USA) for 30-minutes (Barzegarpoor et al., 2020), and then they filled out the NASA Task Load Index (NASA-TLX).

### Statistical Analysis

Statistical analyses were calculated using SPSS software (Version 28.0, SPSS, Inc., Chicago, IL). The Shapiro-Wilk and Mauchly’s Tests were used to assess the normality of the distribution and assumption of sphericity, respectively (P>0.05). The data for time perception were analyzed using the means of six trials. A 3 testing times (pre-test, post-test, and 6-min follow-up) × 3 conditions (control, mental and physical fatigue) with repeated measures analysis of variance (ANOVA) was conducted to determine significant differences for time perception for each time estimate (5-, 10-, 20-, and 30-seconds) separately (within time estimate analysis). One-way repeated measures were conducted to determine significant differences between testing times (pre-test, post-test, and follow-up). A 3 conditions (control, mental, and physical fatigue) × 4 time estimates (5-, 10-, 20-, and 30-seconds) with repeated measures (ANOVA) was conducted to determine significant differences between time estimates. The mean difference (MD) (measures the absolute difference between the mean value in two groups) has been used for the analysis, and it estimates the amount by which the experimental intervention changes the outcome on average compared with another condition. To analyze body temperature and heart rate, an ANOVA with repeated measures were used for 7 testing times (pre-test, start, 10-, 20-, and 30-minutes, post-test, and follow-up) × 3 conditions (control, physical, and mental). To examine RPE during the intervention, a 4 testing times (start, 10-, 20-, and 30-minutes) × conditions (control, mental, and physical fatigue) ANOVA with repeated measures was used. To analyze the NASA Task Load Index, a one-way repeated measures ANOVA was used for mental and physical demand subscales. If the interactions were significant, the Bonferroni post hoc test was conducted to detect the significant differences between conditions for each test. The effect sizes of each variable were tested using partial eta squared (η_p_ ^2^) (0.01= small effect, 0.06= medium effect, 0.14= large effect). The statistical significance level was set at P<0.05. Cohen’s d effect sizes were calculated for individual post-hoc comparisons with effect sizes as trivial (d = <0.2), small (0.2 - ≤0.5), medium (d = 0.5 - ≤0.8), and large (d = ≥0.8) (Cohen 1988)

## Results

### Time estimates

#### Five seconds

The results of the 5-seconds time estimates revealed a significant main effect for the conditions (F_(1.63,26.11)_ =8.44, p=0.003, η_p_ ^2^=0.346) as well as an interaction of testing times and conditions (F_(4,64)_=10.08, p=0.001, η_p_ ^2^=0.387). However, there was no significant main effect for the testing times (F_(2,32)_=3.20, p=0.054, η_p_ ^2^=0.167). There were no significant differences in the interaction of condition * testing time between conditions at the pre-test, but there was a significant large magnitude, underestimation of time for the physical fatigue condition compared to the mental (MD= −0.706 s, p<0.001, d=1.25) and control (MD= −0.577, p<0.001, d=1.45) conditions at the post-test and the follow-up (mental fatigue (MD= −0.842 s, p<0.001, d=1.59), and control (MD= - 0.698 s, p<0.001, d=1.71)). While physical fatigue demonstrated an underestimation of time at post-test and follow-up compared to mental and control conditions, the underestimation with physical fatigue at post-test (MD= −0.539 s) and follow-up (MD= −0.590 s) was also significantly greater than the pre-test, but there was no significant difference between the post-test and follow-up. Additionally, there were no significant differences between the mental fatigue and control conditions during pre-test, post-test, and follow-up. The significant main effect for conditions showed an underestimation of time in the physical fatigue condition compared to mental fatigue (p<0.011, MD= −0.500 s) and control (p<0.001, MD= −0.469 s) conditions. There were no significant differences between mental fatigue and control conditions (Table 2).

**Table 2.** Means and standard deviations of the time estimates of 5-, 10-, 20-, and 30-seconds from the chronological time at the pre-test, post-test, and follow-up.

| Time estimates | Mental fatigue (M±SD) | Physical fatigue (M±SD) | Control (M±SD) |
| --- | --- | --- | --- |
| ***Deviation from 5-seconds from chronological time*** |  |  |  |
| Pre-test | -0.059 ± 0.416 | -0.011 ± 0.288 | 0.121 ± 0.478 |
| Post-test | 0.155 ± 0.751 | -0.550 ± 0.255 | 0.026 ± 0.500 |
| Follow-up | 0.240 ± 0.677 | -0.601 ± 0.314 | 0.096 ± 0.483 |
| ***Deviation from 10-seconds from chronological time*** |  |  |  |
| Pre-test | 0.360 ± 0.734 | 0.301 ± 0.549 | 0.555 ± 0.927 |
| Post-test | 0.669 ± 1.228 | -0.942 ± 0.350 | 0.423 ± 0.898 |
| Follow-up | 1.061 ± 1.446 | -1.006 ± 0.524 | 0.603 ± 1.040 |
| ***Deviation from 20-seconds from chronological time*** |  |  |  |
| Pre-test | 0.581 ± 1.154 | 0.417 ± 1.136 | 1.014 ± 1.542 |
| Post-test | 1.458 ± 2.219 | -1.958 ± 0.538 | 0.748 ± 1.502 |
| Follow-up | 1.830 ± 2.570 | -1.896 ± 0.898 | 0.748 ± 1.502 |
| ***Deviation from 30-seconds from chronological time*** |  |  |  |
| Pre-test | 0.846 ± 1.938 | 0.452 ± 1.733 | 1.281 ± 2.322 |
| Post-test | 1.815 ± 3.459 | -2.947 ± 0.839 | 0.881 ± 2.042 |
| Follow-up | 2.461 ± 3.790 | -2.756 ± 1.413 | 0.943 ± 2.175 |

#### Ten seconds

There were significant main effects with the 10-seconds time estimates for condition (F_(1.40,22.39)_ =12.57, p=0.001, η_p_^2^=0.440), and testing time (F_(2,32)_=4.75, p=0.016, η_p_^2^=0.229) as well as a significant interaction of testing time and conditions (F_(4,64)_=16.91, p=0.001, η^p^ ^2^=0.514). The interaction of condition * testing time showed that there were no significant differences at the pre-test, but there was a significant, large magnitude, underestimation of time in the physical fatigue condition compared to the mental fatigue (MD= −1.612 s, p<0.001, d=1.78) and control (MD= - 1.366 s, p<0.001, d=2.0) conditions at the post-test as well as with the mental fatigue (MD= −2.067 s, p<0.001, d=1.90) and control conditions (MD= −1.609 s, p<0.001, d=1.95) at follow-up. There were no significant differences between mental fatigue and control conditions at the pre-test, post-test, and follow-up. There was a significant (main effect for conditions) underestimation of time in the physical fatigue condition compared to mental fatigue (p=0.001, MD= −1.246 s) and control conditions (p<0.001, MD= −1.077 s). Furthermore, the main effect for the testing time (F_(2,32)_=65.93, p<0.001, η_p_ ^2^=0.805) showed a significant underestimation of time in the follow-up (MD= −1.308 s) and post-test (MD= −1.244 s) compared to the pre-test (Table 2).

#### Twenty seconds

Furthermore, the 20-seconds time estimates revealed a significant main effect for fatigue condition (F_(1.35,21.64)_ =17.12, p=0.001, η_p_ ^2^=0.517), testing time (F =4.27, p=0.037, η_p_ ^2^=0.211), and as well as for the interaction of testing time and conditions (F_(4,64)_=17.12, p=0.001, η_p_ ^2^=0.519). There was a significant, large magnitude, underestimation of time in the interaction of condition * testing time in physical fatigue compared to mental fatigue (MD= −3.418 s, p<0.001, d=2.11) and control (MD= −2.708 s, p<0.001, d=2.39) at the post-test and mental fatigue (MD= −3.726 s, p<0.001, d=1.93) and control (MD= −2.976 s, p<0.001, d=2.13) conditions in follow-up, but there were no significant interactions of condition * testing time at the pre-test among conditions. The main effect for conditions revealed a significant underestimation of time with the physical fatigue condition compared to mental fatigue (MD= −2.436 s, p<0.001) and control (MD= −2.093 s, p<0.001) conditions, but there were no significant differences between mental fatigue and control conditions. The main effect for the testing time demonstrated a significant underestimation of time in the follow-up (MD= −2.314 s) and post-test (MD= −2.376 s) compared to the pre-test, but there were no significant differences between post-test and follow-up (Table 2).

#### Thirty seconds

Thirty seconds showed a significant main effect of fatigue condition (F_(1.28,20.58)_=15.65, p=0.001, η_p_ ^2^=0.495), testing time (F =5.01, p=0.029, η_p2_ =0.239), and as well as for interaction of testing time and conditions (F_(4,64)_=15.90, p=0.001, η_p_ ^2^=0.499). The interaction of condition * testing time revealed that there were no significant differences between conditions at the pre-test, but there was a significant, large magnitude, underestimation of time with physical fatigue compared to mental fatigue (MD= −4.763 s, d=1.89) and control (MD= −3.829 s, d=2.45) conditions at the post-test and mental fatigue (MD= −5.218 s, d=1.82) and control (MD= −3.700 s, d=2.01) conditions in follow-up. Additionally, there were no significant differences between mental fatigue and control conditions at the pre-test, post-test, or follow-up. The main effect for conditions revealed an underestimation with the physical fatigue compared to mental fatigue (MD= −3.458 s, p<0.001) and control (MD= −2.786 s, p<0.001) conditions, but there were no significant differences between mental fatigue and control conditions. A significant main effect for the testing time showed a significant underestimation of time in the follow-up (MD= −3.208 s) and post-test (MD= −3.399 s) compared to the pre-test, but there were no significant differences between post-test and follow-up (Table 2).

#### Relative (%) Time Changes between each Time Estimate

With the post-test, relative time changes showed a significant main effect of conditions (F_(1.27,20.43)_=16.83, p=0.001, η_p2_=0.513), and testing time (F_(1.36,21.89)_=4.12, p=0.044, η_p_ ^2^=0.205), but there was no significant interaction for testing time and conditions (F_(2.60,41.63)_=0.628, p=0.579, η_p_ ^2^=0.038). The main effect for fatigue condition showed that physical fatigue had a significant relative underestimate of time compared to mental fatigue (MD= −0.158 s, p<0.001,) and control (MD= −0.129 s, p<0.001) conditions. There was no significant relative time change between time estimates (5-, 10-, 20-, and 30-seconds) at the post-test with all conditions combined (Figure 4).

**Figure 4.**
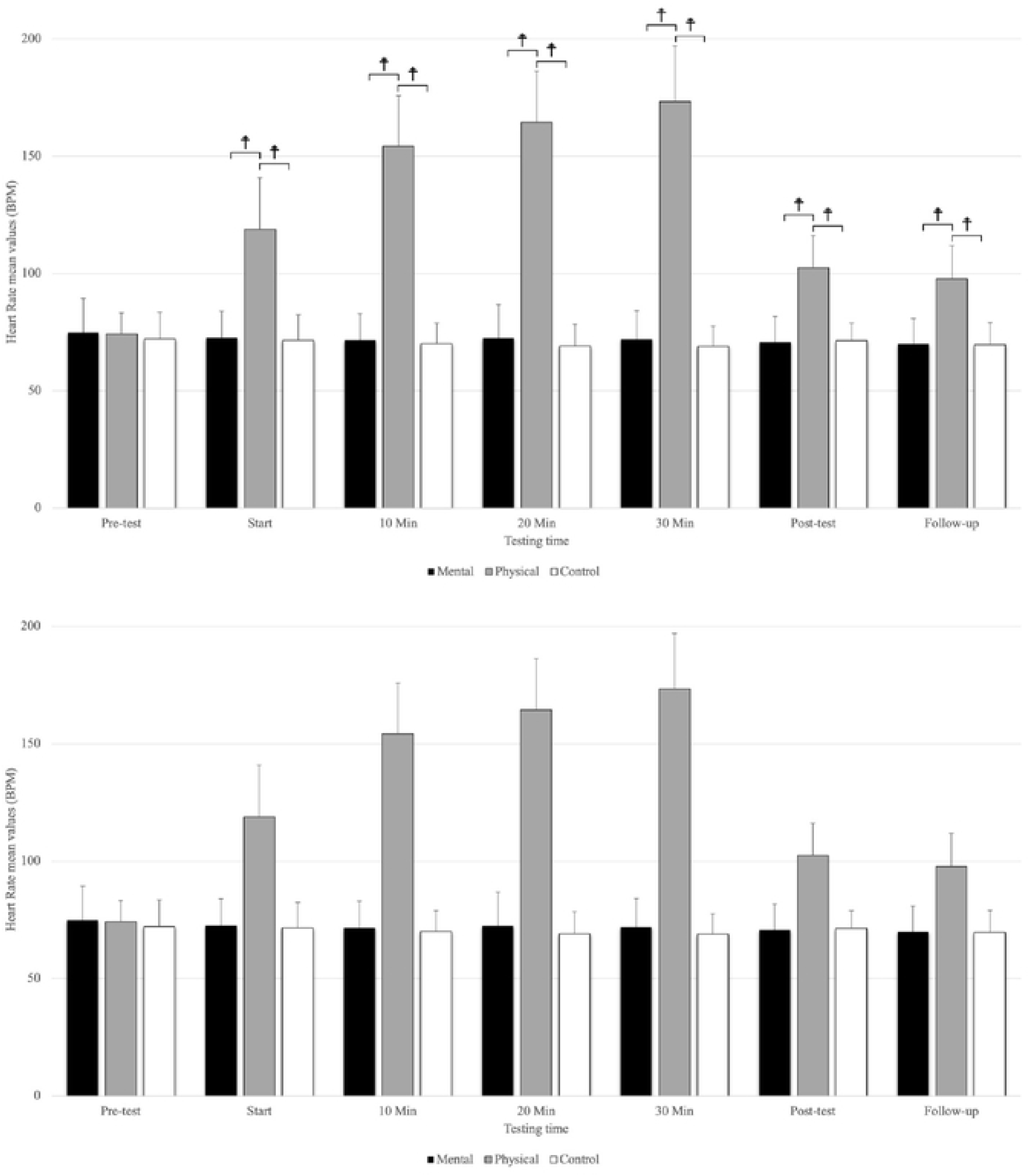
The means and standard deviations of heart rate for the three conditions at the seven testing times. ☨ indicates p < 0.001 compared to the control and mental fatigue condition.

With the follow-up, relative time changes showed a significant main effect of testing time (F_(1.72,27.53)_=3.70, p=0.018, η_p_ ^2^=0.188), conditions (F =18.09, p=0.001, η =0.531), as well as for testing time and conditions (F_(2.32,27.22)_=25.01, p=0.001, η_p_ ^2^=0.610). The main effect for testing time (all conditions combined), revealed relative time changes between 5-, 10-, 20-, and 30-seconds in the follow-up with 10-(MD= 0.040 s, p=0.005) and 20-seconds (MD= 0.044 s, p=0.039) time estimates significantly, relatively higher overestimates than 5-seconds (F_(1.72,27.53)_=3.70, p<0.043, η_p_ ^2^=0.188). There were no significant differences between the 30 seconds with other time estimates (5-, 10-, and 20-seconds) in the follow-up testing period (Figure 5). The main effect for conditions showed that the relative time changes with physical fatigue were significantly underestimated compared to mental fatigue (MD= −0.110 s, p=0.002) and control (MD= −0.125 s, p=0.001) conditions. The interaction of testing time and conditions relative time changes showed that in the mental fatigue, 30-seconds was underestimated compared to the 5-(MD= −0.140 s, p=0.005), 20-(MD= −0.183 s, p<0.001), and 10-seconds (MD= −0.198 s, p<0.001). The results of relative time changes for the physical fatigue revealed that 30-seconds was overestimated compared to the 20-(MD= 0.126 s, p<0.001), 10-(MD= 0.132 s, p<0.001), and 5-(MD= 0.152 s, p<0.001) seconds. Additionally, there was no significant relative time change in the control condition.

**Figure 5.**
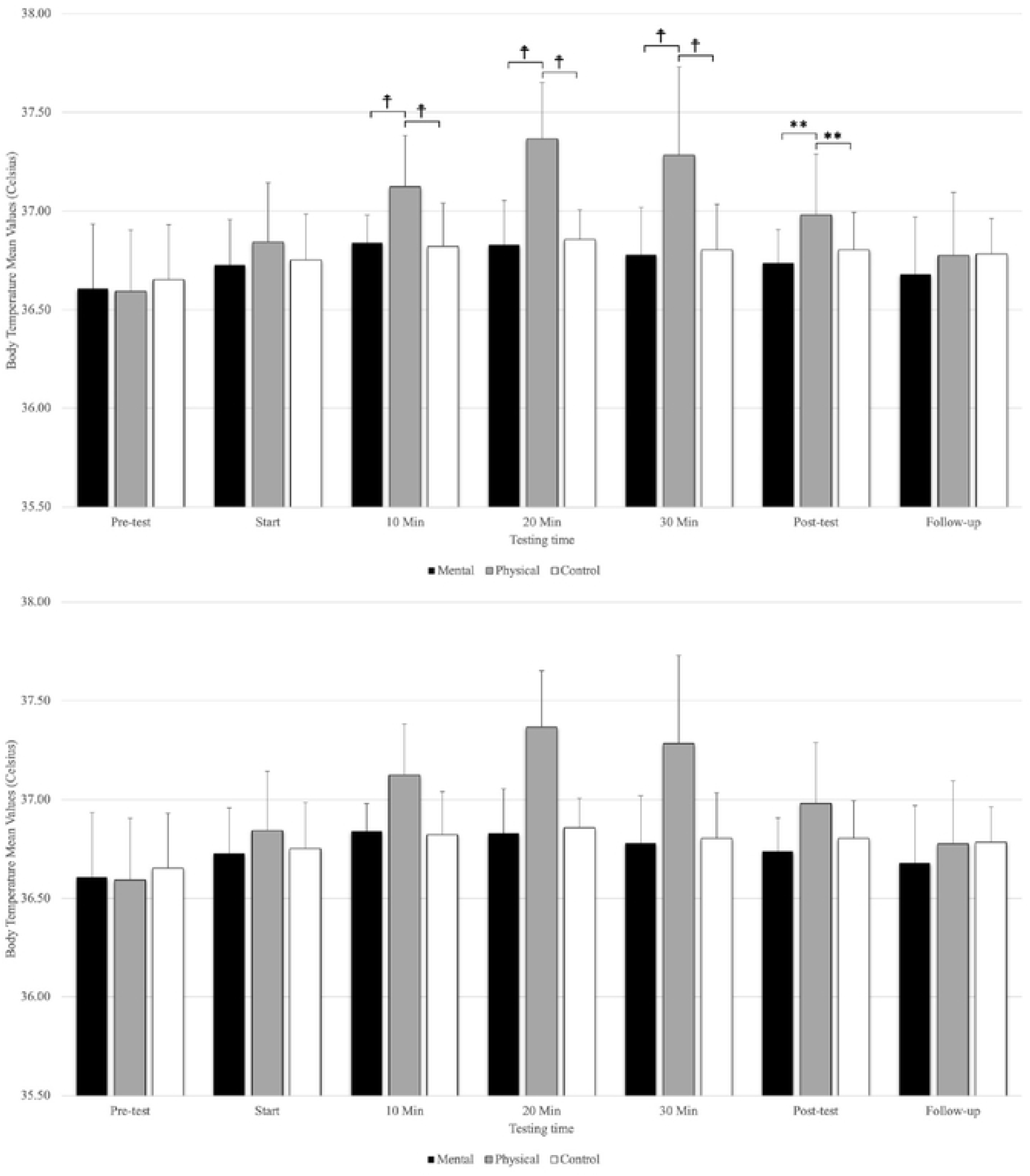
The means and standard deviations of tympanic temperature at the three conditions at the seven testing times. ☨ indicates p < 0.001 compared to the control and mental fatigue condition. ** indicates p < 0.01 compared to the control and mental fatigue condition.

### Heart Rate

Analysis revealed significant main effect for the testing time (F_(3.69,59.09)_ =193.12, p=0.001, η_p_ ^2^=0.923) and conditions (F =286.40, p=0.001, η_p_ ^2^=0.947) as well as a significant interaction of testing time and conditions (F_(4.23,67.73)_ =158.18, p=0.001, η_p_ ^2^=0.908). The condition * testing time interaction showed that there were no significant differences between conditions at the pre-test, but physical fatigue condition (p<0.001) had a significantly higher heart rate compared to the mental fatigue and control conditions at the start,10-, 20-, 30-minutes, post-test, and follow-up (Figure 6). Additionally, there were no significant differences between mental fatigue and control conditions at all testing times. The main effect for conditions showed a significantly large magnitude, higher heart rate for physical fatigue than the mental (MD= 54.513, p<0.001) and control (MD= 56.126, p<0.001) conditions. There was no significant difference between mental fatigue and control condition. The main effects for testing time showed a significant difference (p<0.001) between pre-test, start, 10-, 20-, 30-minutes, and follow-up, except for 20-and 30-minutes (p=0.184) and between post-test and follow-up (p=0.207) (Figure 6).

**Figure 6.**
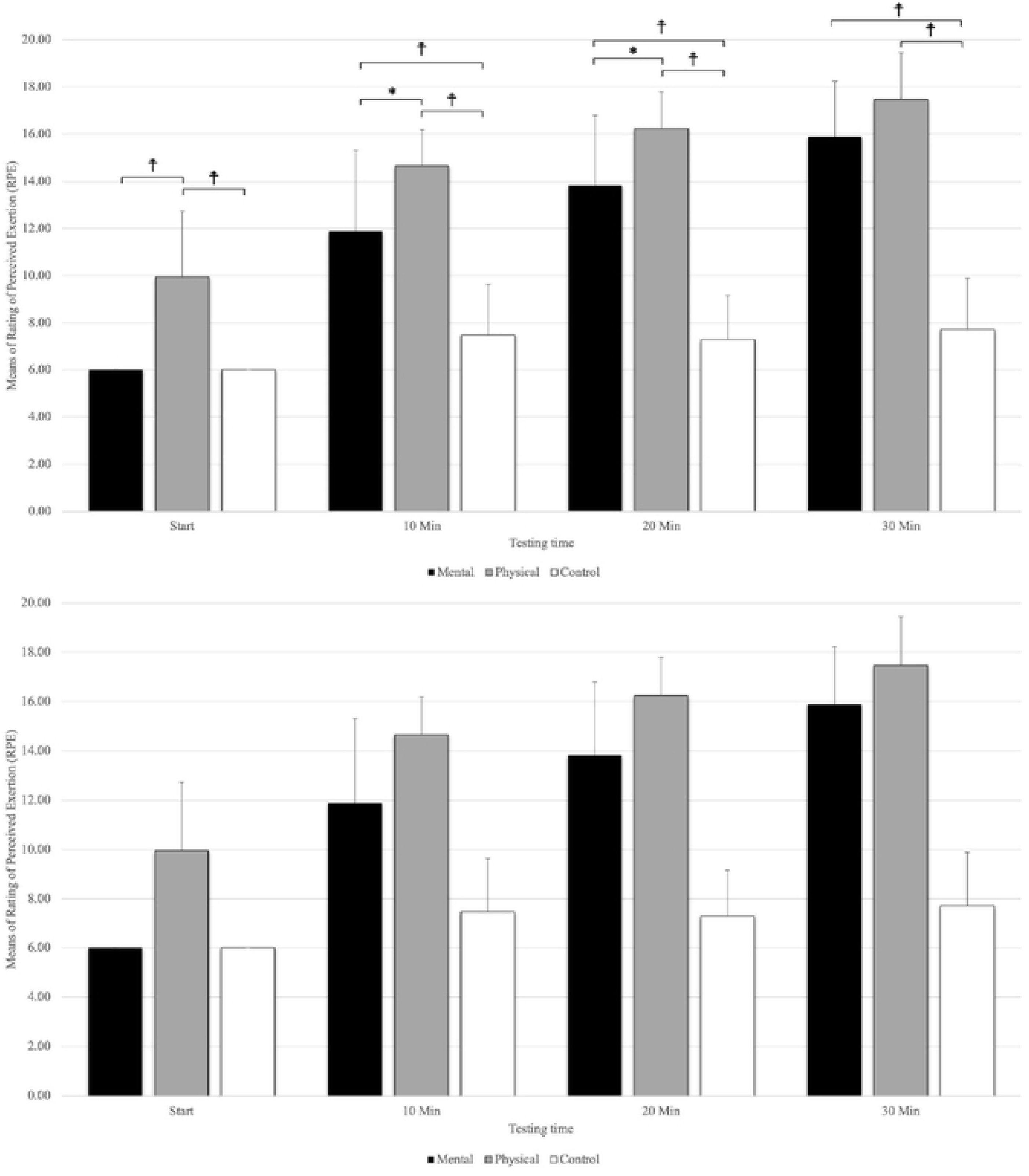
The means and standard deviations of Rating of Perceived Exertion (RPE). ☨ indicates p < 0.001 compared to the control and mental fatigue condition. * indicates p < 0.05 compared to mental fatigue condition.

### Tympanic temperature

A significant main effect was evident for testing time (F_(2.74,43.84)_=27.18, p=0.001, η_p_ ^2^=0.629) and conditions (F_(2,32)_ =13.35, p=0.001, η_p_ ^2^=0.455) as well as the interaction of testing time and conditions (F_(4.70,75.32)_=8.925, p=0.001, η_p_ ^2^=0.358). The interaction of condition * testing time showed that there were no significant differences between conditions at the pre-test and start, but there was significantly elevated tympanic temperature with physical fatigue (p=0<.001) compared to mental fatigue and control conditions at the 10-, 20-, and 30-minutes (Figure 7). There was no significant difference between mental fatigue and control conditions at these testing times (10-, 20-, and 30-minutes). In addition, there was a significantly higher tympanic temperature with physical fatigue (p=0.006) compared to mental fatigue in the post-test. There was no significant difference among conditions at follow-up. The main effect for conditions showed that physical fatigue had significantly, large magnitude, higher tympanic temperature compared to mental fatigue (p=0.001) and control (p=0.008) conditions. Additionally, there were no significant differences between mental fatigue and control conditions. The main effects for a testing time showed a significantly higher tympanic temperature difference in pre-test (p<.001) compared to (start, 10-, 20-, 30-minutes, and post-test), start (pre-test, 10-, 20-, and 30-minutes), 10-minutes (pre-test, start, 20-minutes, and follow-up), 20-minutes (pre-test, 10-, 20-minutes, post-test, and follow-up), 30-minutes (pre-test, start, post-test, and follow-up), post-test (pre-test, 20-, and 30-minutes), and follow-up (10-, 20-, and 30-minutes) (Figure 7).

### Rating of Perceived Exertion (RPE)

A significant main effect was evident for testing times (F_(1.78,28.51)_ =170.51, p=0.001, η_p_ ^2^=0.914) and conditions (F_(2,32)_ =106.80, p=0.001, η_p_ ^2^=0.870) as well as the interaction of testing times and conditions (F_(3.04,48.63)_=24.72, p=0.001, η_p_ ^2^=0.607). The interaction of condition * testing time showed that physical fatigue (p<0.001) had a higher RPE than mental fatigue and control conditions at the start. Physical fatigue (p=0.021) also had a higher RPE than mental fatigue, and mental fatigue (p<0.001) had a higher RPE than control conditions at 10- and 20-minutes (Figure 8). The results revealed that physical fatigue and mental fatigue (p<0.001) had a higher RPE than control conditions, but there were no significant differences between physical fatigue and mental fatigue (p=0.157) at 30-minutes. The main effect for the fatigue condition showed significantly higher RPE scores for physical fatigue versus mental fatigue and control conditions (p<0.001). The main effects for testing time showed a significant difference between RPE (p<.001) the start, 10-, 20-, to 30-minutes (Figure 8).

### The NASA Task Load Index

#### Mental demand

The one-way repeated measure ANOVA (F_(2,32)_=41.87, p<0.001, η_p_ ^2^=0.724) revealed that the mental fatigue condition had a large magnitude, mental demand compared to physical fatigue and control conditions. Additionally, there were no significant differences between physical fatigue and control conditions (Table 3).

**Table 3.** The means and standard deviations (SD) for the mental and physical demand.

| Conditions | Mental Demand<br>Mean ± SD | Physical Demand<br>Mean ± SD |
| --- | --- | --- |
| Mental fatigue | 80±18.28 | 14.41±14.45 |
| Physical fatigue | 34.11±20.63 | 81.17±14.09 |
| Control | 23.23±25.18 | 11.67±17.31 |

#### Physical demand

Physical fatigue had a large magnitude, significant (F_(1.23,19.69)_=167.241, p<0.001, η_p_ ^2^=0.913), and higher physical demand compared to mental fatigue and control conditions, but there were no significant differences between mental fatigue and control conditions (Table 3).

## Discussion

To the best of our knowledge, this is the first study to compare the impacts of mental and physical fatigue on the perception of time. The major findings of this research revealed that participants subjected to physical fatigue exhibited a significant underestimation of time intervals during the post-test and follow-up when compared to those in the mental fatigue and control conditions. In addition, physical fatigue had a significantly higher relative (%) underestimation of time change in comparison to mental fatigue and control conditions. Moreover, physical fatigue induced significantly higher tympanic temperatures and heart rates during the intervention and post-test compared to the mental fatigue and control conditions. The NASA Task Load Index demonstrated the efficacy of both the physical and mental fatigue protocols in inducing states of physical and mental fatigue.

The underestimation of time at 5-, 10-, 20- and 30-s with the physical fatigue condition were in line with the hypothesis predicting significant time underestimations (estimated time was shorter than chronological time) compared to the mental fatigue and control conditions. Moreover, the results for the physical fatigue (underestimation of time) were congruent with Graham et al. (2023), as their results showed an underestimation of time in all three exercise conditions (30-seconds of knee extensors 100%, 60% and 10% of maximum voluntary isometric contraction) with all time estimates (5-, 10-, 20-, and 30-s) compared to the control condition. In addition, the findings of the present study were generally consistent with Gardner et al. (2023), who revealed that maximal contractions induced significantly greater time underestimations at 5-, 20-, and 30-s than control condition. Moreover, their study showed that submaximal (60% of maximal voluntary isometric contractions) contractions also contributed to time underestimation at 30-seconds. Furthermore, Edwards and McCormick (2017) utilized cycling wherein participants were asked to estimate the completion of 25%, 50%, 75%, and 100% of the trial duration under various RPE conditions. Notably, they observed that at the 75% and 100% intervals, time estimates for the RPE 20 condition, representing maximal exertion, exhibited the shortest durations when compared to those of RPE 11 (light intensity) and RPE 15 (moderate intensity). Additionally, the participants also completed a rowing task, wherein they found similar intensity-dependent results (Edwards and McCormick, 2017). Similarly, the RPE findings indicated that participants, upon the end of the physical fatigue intervention, reported an average RPE score of 17. This finding was aligned with the Edwards and McCormick (2017) and suggested that the perceived level of exertion experienced during the physical fatigue condition might be an indicator of underestimation of time in the physical fatigue condition.

In some studies, it has been suggested that an increase in body temperature affects temporal perception (Piéron, 1923; van Maanen et al., 2019). Brinnel and Cabanac (1989), suggested that tympanic temperature as measured in the present study, when measured accurately, is a good index of core temperature and that its variations may reflect variations in brain temperature. Two studies showed that core temperature increased (with running in a warm, humid environment) corresponding to an underestimation of time (Tamm et al., 2014, 2015). Similarly, in the present study, the tympanic temperature was significantly higher in the physical fatigue condition compared to mental fatigue and control during the intervention and the post-test. This finding diverges from the Graham et al. (2023) study, who reported an absence of significant increase in tympanic temperature. Similarly, Gardner et al. (2023) documented that tympanic temperature remained unaffected by the contraction intensities. One possible reason for the observed disparity in outcomes between the present study and the prior investigation lies in the dissimilarities in methodological approaches. Notably, they employed isometric contraction as the primary exercise modality, while we opted for a 30-minute cycling at 65% PPO, which induced a greater rise in tympanic temperature compared to Graham et al. (2023), and Gardner et al. (2023) studies. Moreover, it is pertinent to acknowledge that the duration of their experimental protocol was comparatively shorter than ours, which may have further contributed to differences in physiological reactions and subsequent findings of the two studies.

Another notable finding in this study pertains to the heart rate, which exhibited a significant elevation during the physical fatigue condition at the stages of intervention, post-test, and follow-up in comparison to both the mental fatigue and control conditions. This finding aligns with the findings of Gardner et al. (2023), who reported lower heart rate values for the control condition (75.3 ± 11.6) in contrast to the maximal (92.5 ± 13.9), 60% submaximal (92.2 ± 14.4), or distraction (90.5 ± 14.7) conditions. Similarly, the results obtained by Graham et al. (2023) were consistent with our study, as they demonstrated that the control condition exhibited lower heart rate values (beats per minute) (74.6 ± 10.6) compared to the maximal (91.6 ± 12.4), 60% MVIC (92.5 ± 13.8), or 10% MVIC (90.7 ± 13.5) conditions.

Contrary to our initial hypothesis, the results of our analysis in the mental conditions did not align with our hypothesis, as participants did not exhibit a tendency to underestimate perception of time in this condition. The ideal task duration to induce mental fatigue in young adults is currently unclear in the literature, thus it is not known how long it takes for a change in task performance to become significant. Previous studies have employed tasks that continue for several hours. However, new research indicates that, 30- to 90-minutes is sufficient to cause mental fatigue (Helm, 2021). Individual differences may also be a significant factor in the onset of mental fatigue and the length of time required to induce it. Evidence from some studies suggests that shorter task durations may be sufficient to cause mental fatigue as significant declines in cognitive function were seen after only 30-minutes (Slimani et al., 2018), 45-minutes (Smith et al., 2019), 60-minutes (Wascher et al., 2014) or 90-minutes (Wang et al., 2016). Vrijkotte et al. (2018) in their studies used 90-minutes of Stroop task to induce mental fatigue. The primary conclusion of this research is that in trained, young, healthy athletes, a large magnitude of mental fatigue as determined by the NASA task load index had no impact on physical or cognitive performance (accuracy and reaction times) during the second exercise bout of the two-bout exercise protocol (Vrijkotte et al., 2018). They found that when no mentally fatiguing task was being conducted, the initial maximal exercise test also increased mental fatigue. The individuals were unable to distinguish between physical and mental fatigue (Vrijkotte et al., 2018). Slimani et al. (2018), showed that performing the Stroop task for 30-minutes successfully induced mental fatigue. Although the NASA-TLX showed that Stroop task induced mental fatigue for the participants, the 1100 trials (approximately 30-minutes) might not have been enough to affect time perception or had a shorter duration of impact after the mental fatigue protocol. Another possible mechanism is that mental fatigue and physical fatigue might have different physiological and psychological mechanism that affect time perceptions differently. Our findings for the physical fatigue condition were not consistent with Tonelli et al. (2022,) study, who investigated the effects of moderate physical activity (cycling) on a temporal estimation task in a group of adult volunteers under three different conditions: (1) baseline, (2) during the physical activity phase, and (3) roughly 15 to 20-minutes later, when participants were seated and returned to a resting heart rate (POST). They discovered that exercise directly alters how people perceive time, causing them to overestimate durations in the millisecond range. Notably, the impact lasted during the POST session, ruling out either the heart rate or cycle rhythmicity as the primary contributors (Tonelli et al., 2022).

It was anticipated that when participants estimated the four successive times (5-, 10-, 20-, and 30-seconds), gradually time variability would increase. Naturally, you would anticipate more time variability as time goes on because minor time estimate errors made early in the trial can become more amplified as time goes on (Graham et al., 2023). However, it was intriguing that the relative results showed that 5-, 10-, and 20-seconds intervals demonstrated relatively higher underestimation of time in the physical fatigue condition compared to the 30-seconds.

The findings that physical fatigue can lengthen an individual’s subjective experience of time can be elucidated from the perspective of the Pacemaker-Accumulator Model (PAM) as posited by Gibbon et al. (1984), Grondin (2010), and Allman and Meck (2012). Specifically, in the physical conditions, participants were cycling at 65% of PPO. This physical exertion induced muscle fatigue and discomfort attributed to factors such as tension, partial blood occlusion, and metabolite accumulation, among others. According to Edwards and Polman (2013), this adverse sensation functions as a type of physiological arousal. Arousal has been found to elevate the speed of the pacemaker, resulting in an increased number of pulses accumulated in the accumulator (Gil and Droit-Volet, 2012; Lambourne, 2012). This heightened arousal contributes to a perceived distortion of time, leading to a specific lengthening of perceived time intervals (Gil and Droit-Volet, 2012). Importantly, this time distortion effect exhibits a multiplicative characteristic, wherein the extent of distortion intensifies with longer stimulus durations (Zakay and Block, 1997). Consequently, it is plausible to hypothesize that arousal induced by exercise could engender a time distortion effect (Behm and Carter, 2020; Dormal et al., 2018). In the present study, the heightened state of arousal post-exercise can be attributed to the sustained increase in heart rate and body temperature during the post-test phase. This suggests that the distortion in perception of time persists even after the cycling activity has concluded. The study findings indicate that mental fatigue led to a slight overestimation of time when compared to chronological time; however, the observed difference did not reach statistical significance. Possible reasons for this outcome can be attributed to the limited number of trials employed in the Stroop task, which might not have been sufficient to induce a notable distortion in time perception. Additionally, it is plausible that mental and physical fatigue operate through distinct mechanisms, which could contribute to differential effects on the perception of time. Further investigation and a more comprehensive experimental design are warranted to delve deeper into these intricacies and better comprehend the underlying factors influencing temporal perception in the context of mental and physical fatigue.

### Limitations

This research investigation, akin to any other studies, was not devoid of limitations. One of the hypotheses of the study aimed to compare time estimates between male and female cohorts. However, due to challenges in recruiting an adequate number of female participants, this objective remained unfulfilled. Consequently, the sample primarily consisted of male students engaged in recreational physical activities. An additional limitation of this study pertains to the substantial standard deviations in relation to the mean values, reflecting considerable heterogeneity among the various individual outcomes. Future studies should investigate how mental and physical fatigue might affects perception of time in males and females differently. Additionally, how might time perception be different for endurance exercise above the lactate threshold (e.g., >80-85% max aerobic power)?

## Conclusions

This study’s findings highlight the impact of physical and mental fatigue on participants’ perception of time. Specifically, under the physical fatigue condition, participants underestimated time during the post-test and follow-up, as compared to the mental fatigue and control conditions, across various time intervals (5-, 10-, 20-, and 30-seconds). Moreover, the investigation revealed no significant differences between the mental fatigue and control conditions concerning time estimates. In addition, the results showed that physical fatigue condition demonstrated significantly higher heart rates and body temperatures during both the intervention and post-test, as compared to the mental fatigue and control conditions. Furthermore, participants reported significantly higher RPE under the physical fatigue condition compared to the mental fatigue and control conditions. Additionally, mental demand was significantly higher in the mental fatigue condition than in the physical fatigue and control conditions. The physical demand was significantly greater in the physical fatigue condition relative to both the mental fatigue and control conditions. Overall, these findings contribute valuable insights to the expanding body of research on the relationship between exercise-induced fatigue and time perception.

The present study suggests that individuals engaged in physically demanding activities, such as sports, drivers, and work settings, among others may experience alterations in time perception due to the influence of physical fatigue. Accordingly, it is recommended that these individuals engage in deliberate exercises aimed at enhancing their time perception abilities during periods of physical fatigue. Such practices are hypothesized to facilitate the development of an enhanced sense of timing under physically demanding conditions.

Furthermore, there is another practical aspect to consider: how can we offer feedback or modify exercise routines for individuals who perceive themselves as having limited time available for improvement in order to enhance adherence? This question holds significant relevance, particularly for the general populace that may not find exercise enjoyable. Moreover, the prospective time estimation ratio could also have a notable influence on endurance athletes or time restricted athletes (e.g., tennis, basketball, North American football) who require precise pacing or timing. This factor carries substantial implications, as an athlete who underestimates the time may perform too slowly, jeopardizing their chances of winning a race, whereas an overestimation of time might lead them to push too hard and experience fatigue.

## References

Allman MJ and Meck WH. Pathophysiological distortions in time perception and timed performance. Brain. 2012;135: 656–677. 10.1093/brain/awr210

Barzegarpoor H, Amoozi H, Rajabi H, Button D, Fayazmilani R. The effects of performing mental exertion during cycling exercise on fatigue indices. Intern J Sports Med. 2020; 41: 846–857. 10.1055/a-1179-8326

Behm DG, Carter TB. Effect of exercise-related factors on the perception of time. Front Physiol. 2020;11: 770–775. 10.3389/fphys.2020.00770

Borg, G. Borg’s Perceived Exertion and Pain Scales. 1998; Human Kinetics Publishers.

Bunnag A. The concept of time in philosophy: A comparative study between Theravada Buddhist and Henri Bergson’s concept of time from Thai philosophers’ perspectives. Kasetsart J Social Sci. 2019;40: 179–185. 10.1016/j.kjss.2017.07.007

Colligan L, Potts HW, Finn CT, Sinkin RA. Cognitive workload changes for nurses transitioning from a legacy system with paper documentation to a commercial electronic health record. Intern J Med Informatics 2015;84: 469–476. 10.1016/j.ijmedinf.2015.03.003

Dormal V, Heeren A, Pesenti M., Maurage P. Time perception is not for the faint-hearted? Physiological arousal does not influence duration categorisation. Cognitive Processing 2018;19: 399–409. 10.1007/s10339-017-0852-3

Eagleman DM. Distortions of time during rapid eye movements. Nature Neurosci. 2005;8: 850– 851. 10.1038/nn0705-850

Edwards, M, McCormick A. Time perception, pacing and exercise intensity: maximal exercise distorts the perception of time. Physiol Behav. 2017;180: 98–102. 10.1016/j.physbeh.2017.08.009

Gardner, HR, Konrad A, Alizadeh S, Graham A, Behm DG. Temporal perception is distorted by submaximal and maximal isometric contractions of the knee extensors in young healthy males and females. Front. Sports Active Living 2023;5: 10.3389/fspor.2023.1185480

Gibbon J, Church RM, Meck WH. Scalar timing in memory. Ann NY Acad Sci. 1984;423: 52–77. 10.1111/j.1749-6632.1984.tb23417.x

Gil S., Droit-Volet S. Emotional time distortions: the fundamental role of arousal. Cognition Emotion 2012;26: 847–862. 10.1080/02699931.2011.625401

González-Alonso J, Teller C, Andersen SL, Jensen FB, Hyldig T, Nielsen B. Influence of body temperature on the development of fatigue during prolonged exercise in the heat. J Appl Physiol.1999;86: 1032–1039. 10.1152/jappl.1999.86.3.1032

Graham AP, Gardner H, Chaabene H, Talpey S, Alizadeh S, Behm DG. Maximal and Submaximal Intensity Isometric Knee Extensions Induce an Underestimation of Time Estimates with Both Younger And Older Adults: A Randomized Crossover Trial. J Sports Sci Med. 2023;22: 405–415. 10.52082/jssm.2023.406

Grondin S. Timing and time perception: A review of recent behavioral and neuroscience findings and theoretical directions. Attention Perception Psychophysics 2010;72(3): 561–582. 10.3758/APP.72.3.561

Hart SG, Staveland LE. Development of NASA-TLX (Task Load Index): Results of empirical and theoretical research. In: Advances in Psychology, Elsevier Publishers. 1988;139–183. 10.1016/S0166-4115(08)62386-9

Helm AF. Mental Fatigue: Examining Cognitive Performance and Driving Behavior in Young Adults. University of Massachusetts Doctoral Dissertation 2021; 10.7275/20360886

Iannetta D, Zhang J, Murias JM, Aboodarda SJ. Neuromuscular and perceptual mechanisms of fatigue accompanying task failure in response to moderate-, heavy-, severe-, and extreme-intensity cycling. J Appl Physiol. 2022;133: 323–334. 10.1152/japplphysiol.00764.2021

Ivry RB, Keele S, Diener H. Dissociation of the lateral and medial cerebellum in movement timing and movement execution. Exper Brain Res. 1988;73: 167–180. 10.1007/BF00279670

Jia B, Zhang Z, Feng T. Sports experts’ unique perception of time duration based on the processing principle of an integrated model of timing. PeerJ 2020;8: e8707. 10.7717/peerj.8707

Job RS, Dalziel J. Defining fatigue as a condition of the organism and distinguishing it from habituation, adaptation, and boredom. In: Stress, Workload, and Fatigue. CRC Press. 2000;466–476. 10.1201/b12791-3.2

Kennedy A, Hug F, Sveistrup H, Guével A. Fatiguing handgrip exercise alters maximal force-generating capacity of plantar-flexors. Eur J Appl Physiol. 2013;113: 559–566. 10.1007/s00421-012-2462-1

Krüger RL, Aboodarda SJ, Jaimes LM, Samozino P, Millet GY. Cycling performed on an innovative ergometer at different intensities–durations in men: neuromuscular fatigue and recovery kinetics. Appl Physiol Nutr Metabol. 2019;44: 1320–1328. 10.1139/apnm-2018-0858

Lambourne K. The effects of acute exercise on temporal generalization. Quart J Exper Psychol 2012;65: 526–540. 10.1080/17470218.2011.605959

van Maanen L, Van der Mijn R, van Beurden MH, Roijendijk LM, Kingma BR, Miletić S, et al. Core body temperature speeds up temporal processing and choice behavior under deadlines. Scientific Reports 2019;9: 10053. 10.1038/s41598-019-46073-3

Matell MS, Meck WH. Cortico-striatal circuits and interval timing: coincidence detection of oscillatory processes. Cognitive Brain Research 2004;21: 139–170. 10.1016/j.cogbrainres.2004.06.012

Meck WH. Selective adjustment of the speed of internal clock and memory processes. J Exper Psychol: Animal Behavior Processes 1983;9: 171–176. 10.1037/0097-7403.9.2.171

Meck WH. Neuropsychology of timing and time perception. Brain Cognition 2005;58: 1–8. 10.1016/j.bandc.2004.09.004

Meck WH, Church RM. A mode control model of counting and timing processes. J Exper Psychol: Animal Behavior Processes 1983;9: 320. 10.1037/0097-7403.9.3.320

Mehta RK, Parasuraman R. Effects of mental fatigue on the development of physical fatigue: a neuroergonomic approach. Human Factors 2014;56: 645–656. 10.1177/0018720813507279

Merchant H, Harrington DL, Meck WH. Neural basis of the perception and estimation of time. Ann Reviews Neurosci. 2013;36: 313–336. 10.1146/annurev-neuro-062012-170349

Nobrega AC, O’Leary D, Silva BM, Marongiu E, Piepoli MF, Crisafulli A. Neural regulation of cardiovascular response to exercise: role of central command and peripheral afferents. BioMed Research Intern. 2014; 10.1155/2014/478965

Piéron H. I. Les Problèmes psycho-physiologiques de la perception du temps. L’Année Psychologique 1923; 24: 1–25. 10.3406/psy.1923.4475

Place N, Bruton JD, Westerblad H. Mechanisms of fatigue induced by isometric contractions in exercising humans and in mouse isolated single muscle fibres. Clin Exper Pharmacol Physiol 2009;36: 334–339. 10.1111/j.1440-1681.2008.05021.x

Slimani M, Znazen H, Bragazzi NL, Zguira MS, Tod D. The effect of mental fatigue on cognitive and aerobic performance in adolescent active endurance athletes: insights from a randomized counterbalanced, cross-over trial. J Clin Med 2018;7: 510–517. 10.3390/jcm7120510

Smith MR, Cha R, Nguyen HT, Marcora SM, Coutts AJ. Comparing the effects of three cognitive tasks on indicators of mental fatigue. J Psychol 2019;153: 759–783. 10.1080/00223980.2019.1611530

Smith MR, Coutts AJ, Merlini M, Deprez D, Lenoir M, Marcora SM. Mental fatigue impairs soccer-specific physical and technical performance. Med Sci Sports Exerc. 2016;48: 267–76. 10.1249/MSS.0000000000000762

Stroop JR. Studies of interference in serial verbal reactions. J Exper Psychol. 1935;18: 643. 10.1037/h0054651

Stoet G. PsyToolkit: A software package for programming psychological experiments using Linux. Behavior Res Methods 2010;42: 1096–1104. 10.3758/BRM.42.4.1096

Stoet G. PsyToolkit: A novel web-based method for running online questionnaires and reaction-time experiments. Teaching of Psychol. 2017;44: 24–31. 10.1177/0098628316677643

Tamm M, Jakobson A, Havik M, Burk A, Timpmann S, Allik J, et al. The compression of perceived time in a hot environment depends on physiological and psychological factors. Quart J Exper Psychol 2014;67: 197–208. 10.1080/17470218.2013.804849

Tamm M, Jakobson A, Havik M, Timpmann S, Burk A, Ööpik V, et al. Effects of heat acclimation on time perception. Intern J Psychophysiol. 2015;95: 261–269. 10.1016/j.ijpsycho.2014.11.004

Tonelli A, Lunghi C, Gori M. Moderate physical activity alters the estimation of time, but not space. Front Psychol. 2022;5470. 10.3389/fpsyg.2022.1004504

Vrijkotte S, Meeusen R, Vandervaeren C, Buyse L, Van Cutsem J, Pattyn N, et al. Mental fatigue and physical and cognitive performance during a 2-bout exercise test. Intern J Sports Physiol Perform.2018;13: 510–516. 10.1123/ijspp.2016-0797

Wang C, Trongnetrpunya A, Samuel IBH, Ding M, Kluger BM. Compensatory neural activity in response to cognitive fatigue. J Neurosci. 2016;36: 3919–3924. 10.1523/JNEUROSCI.3652-15.2016

Wascher E, Rasch B, Sänger J, Hoffmann S, Schneider D, Rinkenauer G, et al. Frontal theta activity reflects distinct aspects of mental fatigue. Biol Psycho.l 2014;96: 57–65. 10.1016/j.biopsycho.2013.11.010

Wittmann M. The inner experience of time. Philos Transactions Royal Society B: Biological Sci. 2009; 364: 1955–1967. 10.1098/rstb.2009.0003

Zakay D, Block RA. Temporal cognition. Current Directions Psychol Sci. 1997;6: 12–16. 10.1111/1467-8721.ep11512604

